# Transcriptomic analysis reveals vector attraction to potato virus Y is mediated through temporal regulation of *TERPENE SYNTHASE 1* (*TPS1*)

**DOI:** 10.1101/2024.12.29.630616

**Authors:** Chad T. Nihranz, Prakriti Garg, Junha Shin, Madeleine Dumas, Sunnie Grace McCalla, Sushmita Roy, Clare L. Casteel

## Abstract

Virus-plant dynamics change over time, influencing interactions between plants and insect vectors. However, the signaling pathways and regulators that control these temporal responses remain largely unknown. In this study, we used insect performance and preference bioassays, RNA-Seq, and genetic tools to identify underlying mechanisms mediating temporal variation in plant-virus-vector interactions. We show that settlement and fecundity of the aphid vector, *Myzus persicae*, is increased on potato virus Y (PVY)-infected *Nicotiana benthamiana* plants two weeks after inoculation but not after six weeks. RNA-Seg analysis revealed transcripts related to plant defense and amino acid biosynthesis are upregulated in response to PVY infection and down regulated in response to aphid herbivory, and these patterns changed over time. Based on this analysis we identified a sesquiterpene synthase gene, terpene synthase 1 (*NbTPS1*), that is upregulated early in PVY infection, but not at later infection time points. Using virus-induced gene silencing and transient overexpression in *N. benthamiana* we demonstrate that PVY induction of *NbTPS1* is required for increased aphid attraction to PVY-infected plants in the early stages of infection. Taken together, PVY temporally regulates transcriptional pathways related to plant defense responses and volatile organic compounds that influence aphid vector performance and preference.

## Introduction

Herbivores and pathogens are common threats to plants in nature. Upon perception of a threat, plants initiate signaling cascades that lead to transcriptional reprogramming, resulting in the synthesis of defense-related proteins and secondary metabolites (Tsuda & Somssich, 2015; Ye *et al*., 2021). However, in nature plants must coordinate multiple pathways and defense responses simultaneously as experiencing multiple stresses at once is common (Kliebenstein, 2014; Tsuda & Somssich, 2015). The numerous threats plants face can also vary temporally, which will further require plants to adjust responses dynamically over time (Toruño *et al*., 2016; Wetzel *et al*., 2023). Our understanding of how plants regulate these complex responses in multi-partite interactions is still limited. Thus, advances in this area will be critical in developing effective plant resistance strategies in the future.

Most plant infecting viruses are transmitted by hemipteran insects (e.g., aphids, whiteflies, and leafhoppers) (Whitfield *et al*., 2015). This means that virus infection is most often accompanied by hemipteran feeding, and thus plants must coordinate responses to both challengers in nature. Viruses and their insect vectors have evolved strategies to evade plant defenses or modulate multiple plant responses for their own benefit (Walling, 2008; Wu & Ye, 2020; Ray & Casteel, 2022). Virus-induced changes in plant physiology have also been shown to benefit insect vectors directly through enhanced nutrients and reduced defenses, which increases insect fecundity (Mauck *et al*., 2012; Blanc & Michalakis, 2016). For example, the potyvirus, turnip mosaic virus (TuMV), reduces insect-induced callose deposition and increases the amount of free amino acids in Arabidopsis which enhances the fecundity and survival of *Myzus persicae* (green peach aphid) vectors (Casteel et al., 2014). While the many of the mechanism’s viruses use to manipulate host physiology and enhance their own performance are well understood, the mechanisms underlying virus-induced changes in vector performance are still largely unknown.

Virus-infected plants are frequently more attractive to insect vectors. For example, plants infected with viruses can be more visually attractive to insect vectors (Diener, 1963; Döring & Chittka, 2007) and can produce altered profiles of volatile organic compounds (VOCs) that are used as olfactory cues by insects for host finding (Grunseich *et al*., 2020). Virus impacts on plant physiology and vector-plant interactions can change over the course of infection (Maule *et al*., 2002; Legarrea *et al*., 2015). For instance, rice dwarf virus (RDV) induces the emission of *E*-β-caryophyllene and 2-heptanol in rice (*Oryza sativa*), which differentially affects the behavior of the green rice leafhopper (*Nephotettix cincticeps*) depending on the infection time point (Chang *et al*., 2021, 2023), and aphid emigration changes during disease progression in potato leaf roll virus (PLRV)-infected potato (*Solanum tuberosum* L.), and this was associated with changes in VOC emissions (Werner *et al*., 2009). Despite recent advances, the underlying transcriptional networks that mediate viral impacts on plant-vector interactions are still poorly understood. Despite this, most mechanistic studies of plant-virus-vector interactions have focused on a single time point during infection.

In this study we aimed to develop systems level understanding of how plants respond to multiple threats dynamically over time and uncover the underlying mechanisms using potato virus Y (PVY) and *M. persicae* as model biotic challengers. Previous studies using PVY have shown aphid vectors prefer to settle on PVY-infected potatoes at both early (Bak *et al*., 2019) and late (Srinivasan & Alvarez, 2007) stages of infection however, another study found aphid preference for PVY-infected tobacco (*Nicotiana tabacum* L.) changes over the course of infection (Liu *et al*., 2019). Here we conducted aphid performance and preference bioassays with *Nicotiana benthamiana* plants that had been infected with PVY for two or six weeks. We show *M. persicae* settlement and fecundity is increased on infected plants two weeks after inoculation but not after six weeks. To identify transcriptional pathways differentially regulated by virus infection and aphid herbivory over time RNA-seq was performed with k-means and spectral clustering. Using qRT-PCR, virus-induced gene silencing (VIGS) and transient overexpression, we demonstrate a sesquiterpene synthase gene is differentially regulated over the course of infection, and this mediates vector attraction to virus-infected plants.

## Materials and Methods

### Plants, insects, and virus

*N. benthamiana*, potato virus Y isolate O (PVY^O^), and the PVY vector *M. persicae* (Sulzer) were propagated as previously described (Casteel *et al*. 2014; Bak *et al*. 2019). For mock-inoculated and PVY-infected plants, two leaves of three-week-old *N. benthamiana* were rub inoculated with healthy (mock control) or PVY-infected tissue in a phosphate buffer in a 1 g: 2 mL ratio as previously described (Bak *et al*., 2017). Ten days after inoculation infection was determined by RT-PCR and primers specific to the PVY coat protein as described below (Table S1). Plants were used for experiments at two- and six-weeks post-inoculation (2 wpi, 6 wpi).

### Aphid fecundity bioassays

To assess *M. persicae* performance on early- and late-stage PVY-infected *N. benthamiana*, we conducted fecundity assays as in (Bak *et al*., 2017) using mock-inoculated and PVY-infected plants at 2 wpi and 6 wpi. Briefly, one adult *M. persicae* aphid was placed in a clip cage on the underside of a leaf from each treatment and allowed to produce nymphs. The next day all but one nymph (foundress) was removed from each cage. Ten days later the number of aphids produced from the foundress was recorded. At least 6 separate plants were used for each treatment and the entire experiment was repeated at least twice at 2 and 6 wpi. Fecundity bioassays at 2 wpi and 6 wpi were conducted at different times.

### Aphid choice test bioassays

Choice test assays were conducted using single leaves from mock-inoculated and PVY-infected plants at 2 and 6 wpi as in (Patton *et al*., 2020). Briefly, holes into opposite corners of Nunc^TM^ square dishes (24.3 cm x 24.3 cm x 1.8 cm, Thermo Fisher Scientific, Waltham, MA, USA) and the petiole of one developmentally similar leaf from each treatment was placed into the opposite sides of the dish through the holes. Cotton was placed around the petioles to seal the holes, and the top of the dish placed on top to enclose the leaves in the arena. After two hours ten apterous adult aphids from synchronized colonies (previously starved for 4 hours) were placed in the middle of the arena equidistant from the mock and PVY-infected leaves. After 24 hours, the number of aphids that settled on each leaf and the number undecided were recorded. At least 8 separate choices were conducted at each time point and the entire experiment was repeated at least twice at 2 and 6 wpi.

### RNA-seq of early- and late-stage PVY-infected *N. benthamiana*

At 2 and 6 wpi, 20 third-instar *M. persicae* aphids were clip-caged onto the underside of a leaf from six separate plants of each treatment (mock-inoculated or PVY-infected). Empty clip cages were attached to the underside of a developmentally matched leaf for each treatment as ‘no aphid 0 hour’ controls. Tissue was collected immediately into liquid nitrogen from the empty clip-caged leaves at the start of the experiment (0 hours). For aphid feeding treatments tissue was collected 8 and 48 hours after placing aphids from two separate cages for each treatment. This experiment was replicated 6 times at 2 wpi and 6 wpi (6 replicates x 2 inoculation treatments x 3 aphid treatments x 2 infection time points = 72 plants). These time points were chosen based on our previous study (Bak *et al*., 2017) where we observed that virus-mediated effects on aphid performance were dependent on the re-localization of viral proteins within the first 24 hours of aphid feeding.

Total RNA was extracted from each sample individually using the Quick-RNA^TM^ MiniPrep kit (Zymo Research, Irvine, CA, USA). After RNA extraction two samples from each treatment were pooled (6 samples pooled by 2 = 3 pooled samples per treatment) and used for library prep (three ug of total RNA per pooled sample). Library prep and sequencing were performed at Novogene using an Illumina NovaSeq platform with paired-end 150 bp sequencing (Novogene Corporation Inc., Sacramento, CA, USA). Raw reads were filtered and trimmed by the Illumina pipeline (Parkhomchuk *et al*., 2009), resulting in 52-92 million clean reads (Table S2). Reads were mapped to the newly described *N. benthamiana* genome (Nbe_v1; https://nbenthamiana.jp; Kurotani *et al*., 2023) using Hisat2 v2.0.5 (Mortazavi *et al*., 2008). FeatureCounts v1.4.0-p3 was used to count the mapped reads (Liao *et al*., 2014) and fragments per kilobase of transcript sequence per millions base pairs (FPKM) were calculated based on the length of the gene and read counts mapped. Differential expression analysis was performed using the DESeq2R package (1.20.0) (Anders & Huber, 2010; Love *et al*., 2014). For DEG analysis all 2 wpi virus and aphid samples were compared to 2 wpi mock-inoculated samples, while all 6 wpi virus and aphid samples were compared to 6 wpi mock-inoculated samples. P-values were adjusted using the Benjamini and Hochberg’s approach for controlling the false discovery rate. Genes with an adjusted P-value less than or equal to 0.001 and a log fold change greater than the absolute value of 1.5 were assigned as a differentially expressed gene (DEG).

### PCA analysis and correlation of replicates of RNA-seq transcriptome data

We converted the FPKM values into TPM (Transcripts Per kilobase Million) based on the equation 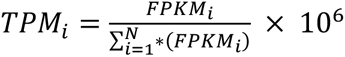, where N is the total number of genes. The data was pre-processed by log-transformation to stabilize variance followed by quantile normalization so that the expression values across different samples are comparable. Finally, we performed zero-means across the samples for centering of each gene value to 0, which resulted in a normalized value matrix of 84570 genes and 36 samples out of 12 conditions (3 replicates). The normalized matrix was leveraged to the Principal Component Analysis (PCA) using MATLAB function ‘pca’ based on Singular Value Decomposition (SVD) algorithm. PCA revealed that triplicate samples from each treatment grouped together (Fig. S1a).

### Transcriptome clustering

The data was pre-processed as described above and then filtered to keep only the genes which had expression values greater than two times the standard deviation in 1% of the samples which resulted in 42968 genes. The expression values of these genes were log-transformed and quantile normalized. Then the three replicates of each of the 12 conditions were collapsed to their means resulting in 12 samples. The data was then zero-meaned across the samples. To identify co-expressed genes in transcriptome clusters, k-means and spectral clustering was performed using k (number of clusters) from 5 to 30 in increments of 5 with Euclidean and Pearson correlation distance metric. Spectral clustering is a graph-based clustering algorithm, which requires us to generate a gene-gene graph. For this, we first constructed a k-nearest neighbour (kNN) distance matrix from the gene expression matrix using Pearson distance and k=20 leveraging ‘knnsearch’ function from MATLAB and made the matrix symmetric requiring the kNN neighbors to be mutual neighbors. Next, the distance matrix was used to calculate the normalized Laplacian eigenvectors corresponding to the smallest k eigenvalues. Finally, to ensure robustness and reliability, k-means was applied with 100 replicates on the selected eigenvalues to identify the clusters. Cluster quality was assessed using Silhouette index (SI). Larger SI values indicate better clustering. Clustering was also performed on non-collapsed data followed by collapsing of replicates to observe the cluster-specific pattern. The final set of clusters across clustering approaches, distance metrics, pre-processing, and number of clusters was selected based on overall extracted patterns, SI, and the ability to recover processes that are expected to be enriched in these clusters. Across our different clustering results, spectral clustering with the Pearson correlation distance metric and k=20 was deemed most favorable and selected for downstream interpretation and analysis.

To interpret our clusters, we used Gene Ontology (GO) process term enrichment using an FDR-controlled Hyper-geometric test. We shortlisted the most important enriched GO terms for the spectral clusters, on-negative matrix factorization (NMF) was done on the negative log of FDR-corrected P-values obtained from the hypergeometric tests using an approach similar to muscari (Lee & Seung, 1999; Shin *et al*., 2021). Briefly, NMF was applied to group the clusters and GO terms in three biclusters. Next, from each of the three GO pathway biclusters, we took the top four GO terms based on their contribution to each factor, followed by a greedy approach where a GO terms was selected only if it was enriched for a spectral cluster which was not previously included based on selected pathways. This enabled us to have a good coverage of both GO terms and spectral clusters. For visualization, we included all non-zero 17 spectral clusters. The key terpenoid and linolenic acid related pathways were also included in the visualization, as there are related to volatile production and plant defense.

### RT-PCR and quantitative RT-PCR

Reverse transcription PCR (RT-PCR) was used to verify infection, and quantitative RT-PCR was used to quantify PVY titer in plants, validate RNA-Seq results, and confirm silencing of *NbTPS1*. Total RNA was extracted as described above. First-strand cDNA was synthesized from 1500ng of RNA using oligo dT (SMART® MMLV, Takara Bio USA, Inc, San Jose, CA, USA) and diluted 1:10 with molecular grade water. Primers were designed to a conserved region of three *N. benthamiana* terpene synthetase sequences (Nbe_v1_s00120g40900, Nbe_v1_s00120g40920, and Nbe_v1_s00120g40940). These sequences were identified in the RNA-seq experiment and overlapped at >99.9% to the *TERPENE SYNTHASE* 1 coding sequence from *N. benthamiana* (*NbTPS1*; GenBank: KF990999.1). Primers were also designed to the PVY coat protein coding sequence and to *ELONGATION FACTOR 1ɑ*, which served as the reference gene (Table S1). All qRT-PCR reactions used SsoAdvance™ Universal SYBR® Green Supermix (BioRad Laboratories, Inc., Hercules, CA, USA) and were run on a CFX384™ Optics Module Real-Time System (BioRad Laboratories, Inc., Hercules, CA, USA). The program had an initial denaturation for 2 min at 94°C followed by 40 cycles of 94°C for 15 sec and 55°C for 30 sec. Relative expression of genes was calculated using the delta-delta Ct method (2^-ΔΔCt^) (Livak & Schmittgen, 2001).

### Virus-induced gene silencing (VIGS) and aphid choice experiments

For VIGS, we amplified 292 bp conserved region of the three *N. benthamiana* sequences (Nbe_v1_s00120g40900, Nbe_v1_s00120g40920, and Nbe_v1_s00120g40940) using gene specific primers (Table S1). Ligation independent cloning was used to insert the sequence into the Tobacco Rattle Virus 2 construct (TRV2-LIC) as in Dong et al. (2007). In brief, amplicon insert and TRV2-LIC vector overhangs were created. The TRV2-LIC vector was linearized and mixed with 50 ng of the amplicon in a 1:1 (v/v) ratio, then transformed into *Escherichia coli* (strain DH5ɑ) and *Agrobacterium tumefaciens* (strain GV3101) (Hofgen & Willmitzer, 1988). A 1:1 (v/v) ratio of *A. tumefaciens* containing either TRV1 and TRV2 or TRV1 and TRV2:*NbTPS1i* silencing construct at a OD_600_ of 1.0 was infiltrated into leaves of 3-week-old *N. benthamiana* using a needleless syringe as previously described (Prakash *et al*., 2023). Plants were either mock-inoculated or infected with PVY as described above before inoculations. Plants were used 14 days later in aphid choice test bioassays as described above. For paired choice tests, aphids were given a choice between leaves of (1) mock-inoculated plants with TRV1+TRV2 (TRV) control and PVY-infected plants with TRV control or (2) mock-inoculated plants with TRV control and PVY-infected plants with TRV1+TRV2:*NbTPS1i* (TRV:*NbTPS1i*) silencing construct.

### Transient overexpression and aphid choice experiments

The *NbTPS1* coding sequence (GenBank: KF990999.1), was synthesized with a Myc tag at the 5’ end and His at the 3’ end, and cloned into the pMDC32 expression vector between the KpnI and SpeI restriction sites by Twist Bioscience (San Francisco, CA). The construct was transformed into *E. coli* (strain DH5ɑ), and transformed colonies were verified by PCR. Successful transformants were used to amplify the construct, then the purified construct from *E. coli* cultures was used to transform *A.* tumefaciens as above. Healthy leaves of 4-week-old *N. benthamiana* were infiltrated with either a pMDC32 empty expression vector (pMDC32 EV) or pMDC32 containing *NbTPS1* (pMDC32:*NbTPS1*) at a OD_600_ of 0.2. Two days later dual choice tests were performed as described above. Overexpression of *NbTPS1* was confirmed by Western blot as previously described (Bera *et al*., 2022) with 1:5000 dilution of Myc tag monoclonal antibody (ThermoFisher Scientific, Waltham, MA, USA) and a 1:10,000 dilution of goat anti-mouse IgG-HRP (Santa Cruz Biotechnology, Dallas, TX, USA). In all agroinfiltrations, the P19 silencing suppressor from Tomato bushy stunt virus (TBSV) was also infiltrated into leaves to enhance expression of VIGS and transient overexpression constructs.

### Statistical analysis

Data analyses were performed using R statistical software (R Development Core Team, 2013) or MATLAB v R2022b Update 6. A Mann-Whitney U test was used to assess the effect of early- and late-stage PVY infection on the number of nymphs produced, as aphid fecundity data did not meet the assumption of normality. Log-likelihood test of independence (‘*GTest*’) was used to analyze all aphid choice tests. Two-way ANOVAs were used to analyze transcript abundance of *NbTPS1* from the RNA-seq data with virus treatment, aphid treatment, and their interaction as fixed effects. Tukey post hoc tests were performed to determine differences between treatments groups. An upset plot (‘*UpSetR*’) was created to visualize DEGs unique to individual aphid treatments and shared among aphid treatments (Conway *et al*., 2017). Heatmaps were created with the ‘*pheatmap*’ package (Klode, 2019). For aphid choice tests using TRV, RT-qPCR data could not be transformed to achieve normality, so Kruskal-Wallis tests were performed to assess PVY coat protein and *NbTPS1* relative expression. Post hoc Dunn’s tests for multiple comparisons with Bonferroni adjustments were performed to determine the differences between treatment groups in these choice tests. Figures were created with ggplot2 in R (Wickham, 2016).

Principal Components Analysis (PCA) for the dataset was done with MATLAB v R2022b Update 6 (see above for details). Gene Ontology enrichment analysis was done using a hypergeometric test followed by a FDR cutoff of 0.05 and further analyzed using MATLAB v R2022b Update 6 NMF implementation via nnmf function with parallel processing (see above for details).

## Results

### PVY increases aphid fecundity and settlement during early stages of infection

To determine how PVY infection affects plant-aphid interactions over time, we assessed aphid fecundity and aphid settlement on mock-inoculated and PVY-infected plants at 2 and 6 wpi. At 2 wpi, the number of aphids was significantly higher on PVY-infected plants compared to mock-inoculated plants (Fig. 1a; W = 398.5, *P* = 0.003), while at 6 wpi, there was no difference in the number of aphids between treatments (Fig. 1a; W = 409, *P* = 0.422). At 2 wpi, more aphids settled on PVY-infected plants compared to mock-inoculated plants (Fig. 1b; G = 7.713, *P* = 0.005). In contrast, at 6 wpi, there was no difference in aphid settlement between treatments (Fig. 1b). PVY titer was significantly greater in PVY-infected plants at 6 wpi compared to 2 wpi (Fig. 1c; F_1,14_ = 25.285, *P* < 0.001), indicating that plants were more infected at later stages but that this increase in viral titer did not influence aphid choice. Taken together, this suggests that aphid vector performance and preference were enhanced by PVY early in infection, but not later stages of infection.

**Fig. 1:**
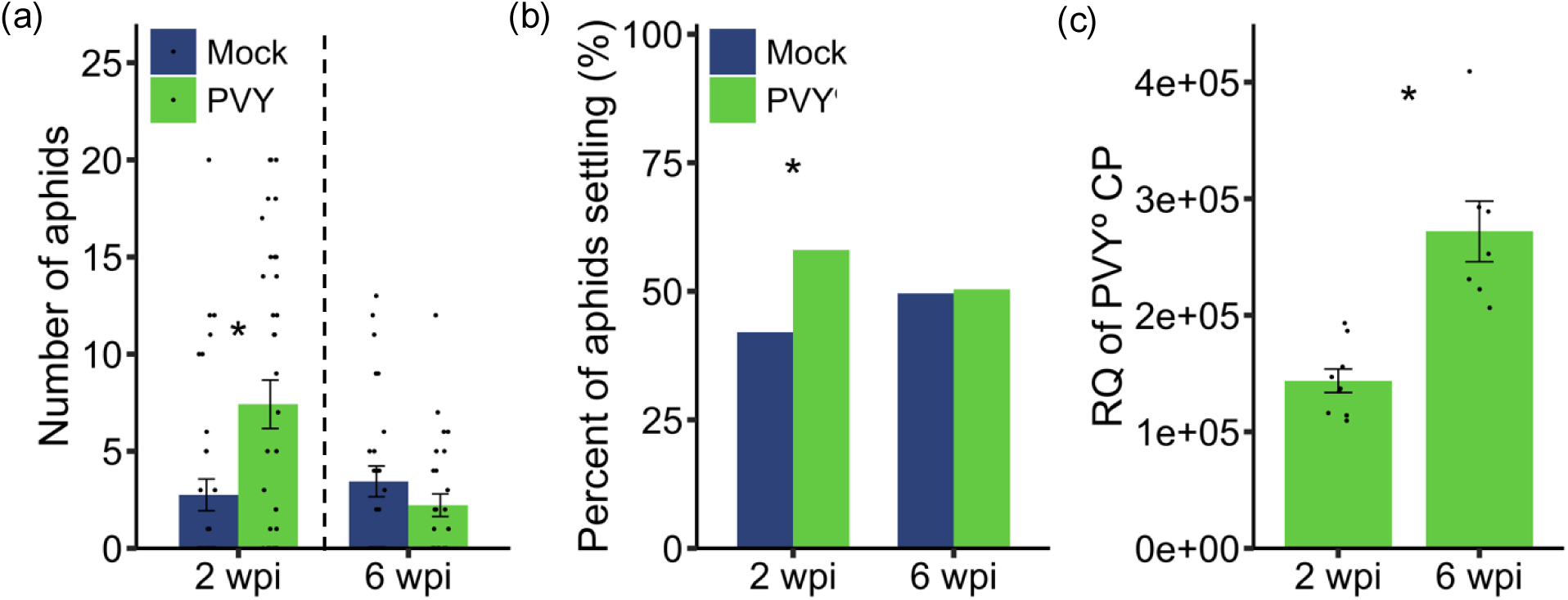
*Myzus persicae* aphids have increased fecundity and settlement on PVY–infected *Nicotiana benthamiana* early in infection but not later in infection. (a) A single *M. persicae* aphid nymph was clip caged on to the underside of leaves of mock-inoculated and PVY-infected *N. benthamiana* at 2- and 6-weeks post inoculation (wpi). Ten days later, the number of aphids in each clip cage was recorded. (b) Adult *M. persicae* apterous aphids were given a choice between leaves of mock-inoculated and PVY-infected *N. benthamiana* plants at 2 wpi and 6 wpi. Twenty-four hours later aphid settlement was assessed. (c) Relative quantification (RQ) of PVY coat protein (CP) in *N. benthamiana* leaves infected with PVY at 2 wpi and 6 wpi relative to mock-inoculated plants of the same age. Asterisks indicate significant differences *P* < 0.01 between mock-inoculated and PVY-infected plants according to a Mann-Whitney U test for (a) (N = 13-24;) and to a Log-likelihood test of independence test for (b) (N = 16). For (c) asterisks indicate significant differences between PVY-infected plants at 2 wpi and 6 wpi (ANOVA; N = 7-9; *, *P* < 0.001).

### PVY infection triggers strong transcriptional responses at early time points that dampen over time

We performed RNA-seq to examine the transcriptional response of plants to early- and late-stage PVY infection and *M. persicae* herbivory. Overall, there were 11,160 differentially expressed genes (DEGs) across all treatments (Fig. 2a-d; Table S3). PVY infection alone resulted in 2893 DEGs at 2 wpi and 608 DEGs at 6 wpi compared to mock-inoculated plants of the same age (Fig. 2a). Of these, only 197 DEGs were shared between the 2 and 6 wpi PVY alone treatment (Fig. 2b). Most of the DEGs were upregulated in response to PVY (Fig. 2a; 2 wpi PVY: 1602 up/1291 down; 6 wpi PVY: 453 up/155 down). These results demonstrate PVY infection triggering strong initial transcriptional response that decreases over time.

**Fig. 2:**
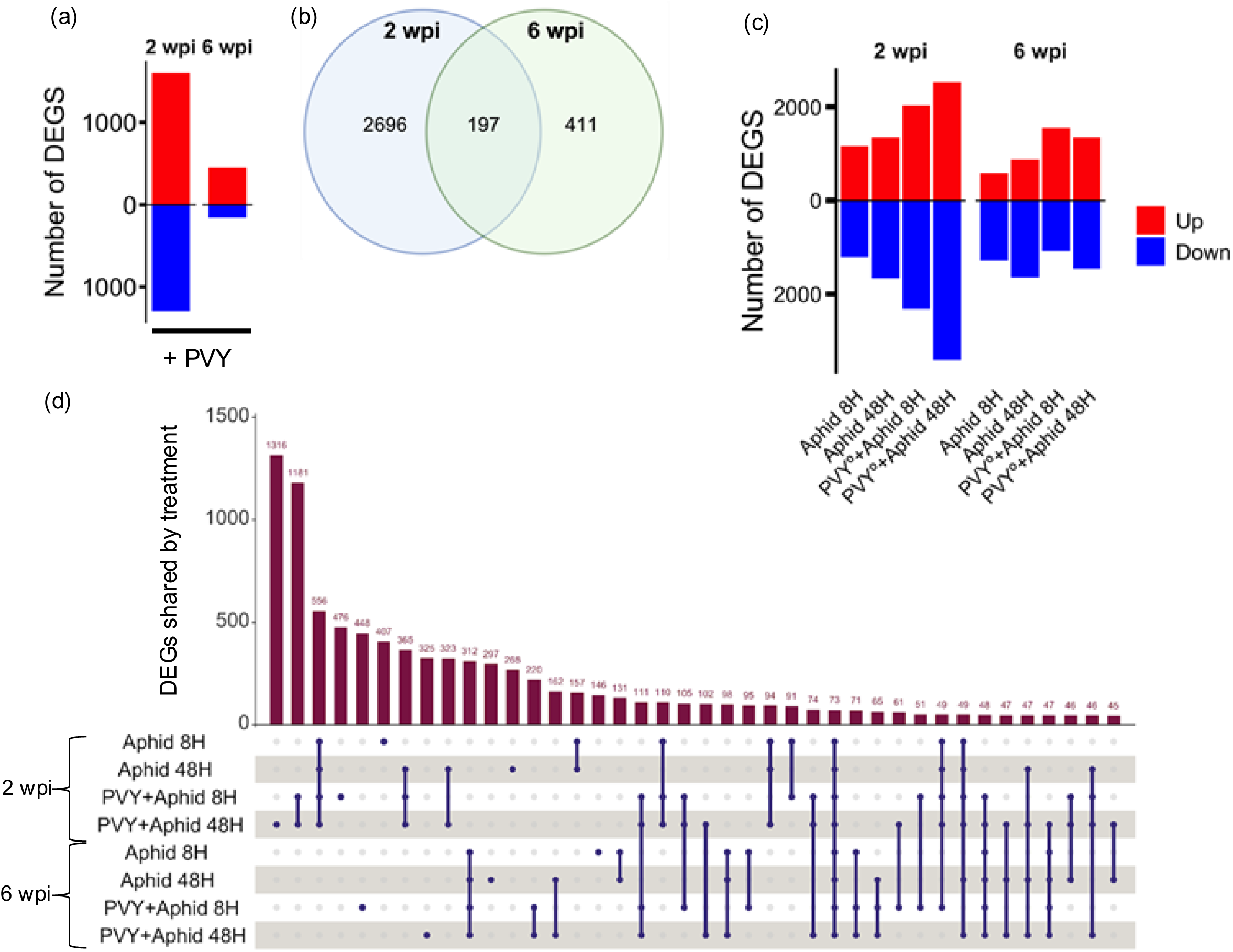
PVY infection triggers strong transcriptional responses at early time points that dampen over time. (a) Number of significantly up-regulated (red) and down-regulated (blue) DEGs in PVY-infected *N. benthamiana* plants at 2- and 6-weeks post-inoculation (wpi). (b) Venn diagram showing number of DEGs unique to and shared by PVY-infected *N. benthamiana* plants at 2 and 6 wpi. (c) Number of significantly up-regulated (red) and down-regulated (blue) DEGs by 8 hours (8H) and 48 hours (48H) of *Myzus persicae* herbivory on mock-inoculated and PVY-infected *N. benthamiana* plants at 2 wpi and 6 wpi. (d) Upset plot illustrating unique and shared DEGs across *M. persicae* herbivory treatments. Each vertical bar (dark red) in the bar graph represents the number of DEGs unique to treatments or shared among treatments. The number of DEGs is indicated above each bar. The matrix grid displays the presence or absence of unique and shared DEGs in each treatment. Single blue circles indicate DEGs unique to that treatment. Connected blue circles indicate shared DEGs between or among treatments. Data in all figures are number of significantly up- and down-regulated DEGs of 6 biological replicates determined by negative bionomical distribution (Wald test). DEGs were included in analysis only if they had a log fold change greater than the absolute value of 1.5 and a P-value less than 0.001. The complete list of DEGs can be found in Supporting Information Table S3. (e)

In contrast to PVY responses, a similar number of *N. benthamiana* genes were differentially regulated at the transcriptional level in response to *M. persicae* feeding in the mock treatments, regardless of herbivory time treatment (8 or 48 hr) or plant age (2 or 6 wpi) (Fig. 2c; Table S3; Mock 2 wpi: 8 hr/2378 & 48 hr/3012 DEGS; Mock 6 wpi: 8 hr/1865 & 48 hr/2525 DEGS). Younger plants that had been infected with PVY and infested with *M. persicae* responded with almost twice as many DEGS as compared to the 6 wpi PVY aphid treatments (Fig. 2c; Table S3; PVY 2 wpi: 8 hr/4353 & 48 hr/5942 DEGS; PVY 6 wpi: 8 hr/2629 & 48 hr/2802 DEGS). Only 157 and 131 DEGs were shared among the 8- and 48-hour herbivory treatments at the 2 and 6 wpi mock treatments respectively, while 1181 and 220 DEG were in common for the 8- and 48-hour herbivory between 2 and 6 wpi PVY treatments (Fig. 2d). Among these only 73 DEGs were shared between all aphid herbivory treatments regardless of infection status or duration of aphid feeding time. Thes genes may represent core plant regulators of plant responses to aphid herbivory. Taken together, these results suggest that plant responses to PVY infection are more dynamic than compared to aphid herbivory.

### Transcripts related to plant defense and amino acid biosynthesis are upregulated in response to PVY infection and down regulated in response to aphid herbivory

To determine the system level patterns of co-expression at the mRNA level across PVY and *M. persicae* treatments, we used spectral clustering and GO enrichment on our complete dataset of ∼43k expressed genes (Fig. 3a,b). Of the 20 clusters identified, 17 were enriched for GO terms and were examined across treatments (Fig. 3a,b; Table S4). Transcripts in cluster C17 (1146 genes) increased in response to PVY infection, with this increase becoming more pronounced throughout the infection cycle. This cluster was enriched in GO terms related to plant defense (biosynthesis of secondary metabolites, plant hormone signal transduction, MAPK signaling, phenylpropanoid biosynthesis, cutin, suberin, and wax biosynthesis, and sesquiterpenoids and triterpenoids biosynthesis) and biosynthesis of numerous amino acids (Fig. 3b; Table S4). Conversely, aphid herbivory downregulated the expression of transcripts in C17. Transcripts in clusters C4 (1866 genes) and C9 (1377 genes) had reduced expression in PVY-infected plants compared to mock-inoculated plants at 2 wpi but not at 6 wpi (Fig. 3b; Table S4). Clusters C4 and C9 were both enriched for biosynthesis of cofactors and secondary metabolites, while C4 was enriched for photosynthesis processes and terpenoid backbone biosynthesis GO terms.

**Fig. 3:**
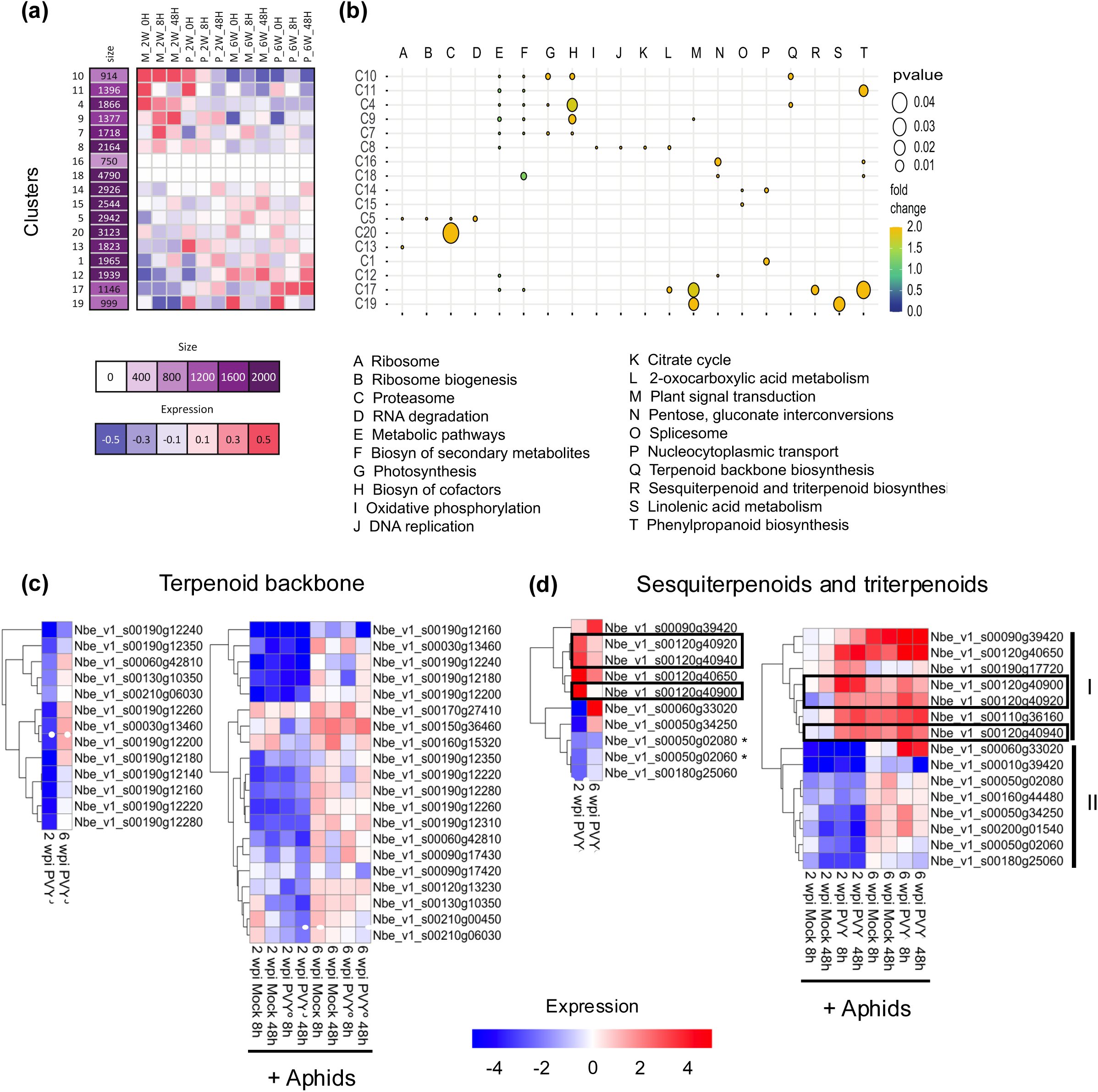
Transcripts related to secondary metabolism are regulated dynamically in response to PVY and aphid herbivory over time. (a) Spectral clustering of transcriptome from mock-inoculated or PVY-infected *N. benthamiana* with and without aphid herbivory. Columns indicate treatments and rows are associated clusters. The first column indicates the “size” or total number of genes co-expressed in each cluster with darker purple colors indicating a greater number of genes in that cluster. Subsequent columns indicate treatments and cells below each treatment show expression values for each cluster. (b) Plot of GO terms enriched in each cluster. Clusters are shown on the on the left side of the plot. Size of circle indicates fold change of GO term associated with that cluster (ranging from 0 to 2.0) and color of circle indicates associated P-value. C, cluster; M, mock-inoculated plants; P, PVY-infected plants; 2W, 2 weeks post inoculation; 6W, 6 weeks post inoculation; 0H, no aphid herbivory; 8H, 8 hours of aphid herbivory; 48H, 48 hours of aphid herbivory. Heatmaps of differentially expressed genes in KEGG pathways related to (c) terpenoid backbone biosynthesis and (d) sesquiterpenoid and triterpenoid biosynthesis in mock-inoculated and PVY-infected *N. benthamiana* plants at 2 wpi and 6 wpi without herbivory and with *M. persicae* aphid herbivory for 8 hours (8h) and 48 hours (48h). Red indicates up-regulation and blue cells indicates down-regulation. Each row represents a different gene within each pathway. Black boxes in (d) indicate terpene synthase genes that were significantly upregulated by PVY early in infection but not at later stage of infection. Asterisks (*) indicate genes that synthesize triterpenoid products.

### PVY and aphid herbivory dynamically regulate transcripts related to sesquiterpenoid and triterpenoid biosynthesis

Because terpenoids are known to influence insect behavior (Boncan *et al*., 2020) and terpenoid backbone, sesquiterpenoid, and triterpenoid terms were enriched among multiple expression clusters (C4, C10, and C17; Fig. 3a,b), we next examined these pathways in more detail using KEGG analysis. KEGG analysis revealed 21 DEGs in the terpenoid backbone pathway were significantly downregulated in response to PVY at 2 wpi relative to the controls, and 13 of these DEGs synthesize immediate precursors of volatile terpenoid products (Fig. 3c). Only 3 DEGs in this pathway were significantly regulated by PVY at 6 wpi compared to controls at the same timepoint, and none of these three synthesized immediate precursors of terpenoid volatile products (Fig. 3c). Of the 44 DEGs in the terpenoid backbone pathway significantly regulated by aphid herbivory, 20 were related to the synthesis of volatile terpenoid products (Fig. 3c). At 2 wpi, these genes were largely downregulated in response to aphid herbivory on mock-inoculated and PVY-infected plants relative to controls at the same time points.

In the sesquiterpenoid and triterpenoid pathway, five DEGs predicted to encode key enzymes involved in the production of plant volatile organic compounds were significantly downregulated by PVY infection at 2 wpi compared to controls at the same time point (Fig. 3d). These include β-amyrin synthase (Nbe_v1_s00050g02060 and Nbe_v1_s00050g02080), which synthesizes the triterpenoid β-amyrin, (3S,6E)-nerolidol synthase (Nbe_v1_s00050g34250), which synthesizes (3S,6E)-nerolidol, α-farnesene synthase (Nbe_v1_s00060g33020), which synthesizes α-farnesene, and (-)-germacrene D synthase (Nbe_v1_s00180g25060), which synthesizes (-)-germacrene D. In contrast, three DEGs that encode 5-epi-aristolochene synthase (Fig 3d boxed genes; Nbe_v1_s00120g40900, Nbe_v1_s00120g40920, and Nbe_v1_s00120g40940) and one DEG that encodes 5-epi-aristolochense 1,3-dihydroxylase (Nbe_v1_s00120g40650), which are involved in the synthesis of 5-epi-aristolochene and capsidiol, respectfully, were significantly upregulated by PVY at 2 wpi compared to controls at the same time point (Fig 3d). At 6 wpi, PVY upregulated the expression of only one DEG in these pathways, another 5-epi-aristolochense 1,3-dihydroxylase (Nbe_v1_s00090g39420), which was not significantly affected at 2 wpi.

We identified two groups of DEGs in the sesquiterpenoid and triterpenoid pathway that were differentially affected by aphid herbivory with and without PVY infection (Fig. 3d). DEGs in group I were upregulated by aphid herbivory on PVY-infected plants at 2 wpi but not on mock-inoculated plants at the same time point. Group I includes (-)-germacrene D synthase (Nbe_v1_s00010g39420), squalene synthase (Nbe_v1_s00190g17720), 5-epiaristolochene 1,3-dihydroxylase (Nbe_v1_s00120g40650), and 5-epi-aristolochene synthases (Nbe_v1_s00110g36160, Nbe_v1_s00120g40900, Nbe_v1_s00120g40920, and Nbe_v1_s00120g40940). At 6 wpi, aphid herbivory largely upregulated the genes in group I regardless of infection status. DEGs in group II were largely downregulated by aphid herbivory on mock-inoculated and PVY-infected plants at 2 wpi and included β-amyrin synthase (Nbe_v1_s00050g02060, Nbe_v1_s00050g02080, and Nbe_v1_s00200g01540), (-)-germacrene D synthase (Nbe_v1_s00180g25060), squalene synthase (Nbe_v1_s00160g44480), α-farnesene synthase (Nbe_v1_s00060g33020), (3S,6E)-nerolidol synthase (Nbe_v1_s00050g34250) and 5-epiaristolochene 1,3-dihydroxylase (Nbe_v1_s00090g39420). These results illustrate that PVY infection and aphid herbivory have complex and dynamic effects on the expression of transcripts related to volatile terpenoid biosynthesis over time.

### PVY induction of terpene synthase 1 (*NbTPS1*) early in infection is required for increased aphid preference

Since 5-epi-aristolochene has been shown to be involved in plant-virus-vector interactions (Li *et al*., 2014) and three genes that encode 5-epi-aristolochene synthase were significantly upregulated by PVY at 2 wpi but not 6 wpi (Fig. 3d; boxed genes in PVY alone heat map), we decided to investigate the biological relevance of these changes in relation to aphid vector behavior. Sequence alignment revealed that these three genes are 99.3% identical to each other and overlapped over 99% with the *NbTPS1* coding sequence (GenBank: KF990999.1). Given sequence similarity, a single set of primers were used to confirm *NbTPS1* transcript changes in the RNA-seq results via RT-qPCR. Consistent with the RNA-Seq results, RT-qPCR validated that *NbTPS1* expression was significantly increased by PVY infection at 2 wpi but not at 6 wpi relative to controls at the same time points (Fig. 4a). *NbTPS1* expression was also significantly increased by aphid herbivory on PVY-infected plants at 2 wpi relative to controls at the same time points (Fig. 4a); however, there was no difference between aphid induction of the transcripts between mock or PVY-infected plants at 6 wpi relative to controls at the same time points (Fig. 4a).

**Fig. 4:**
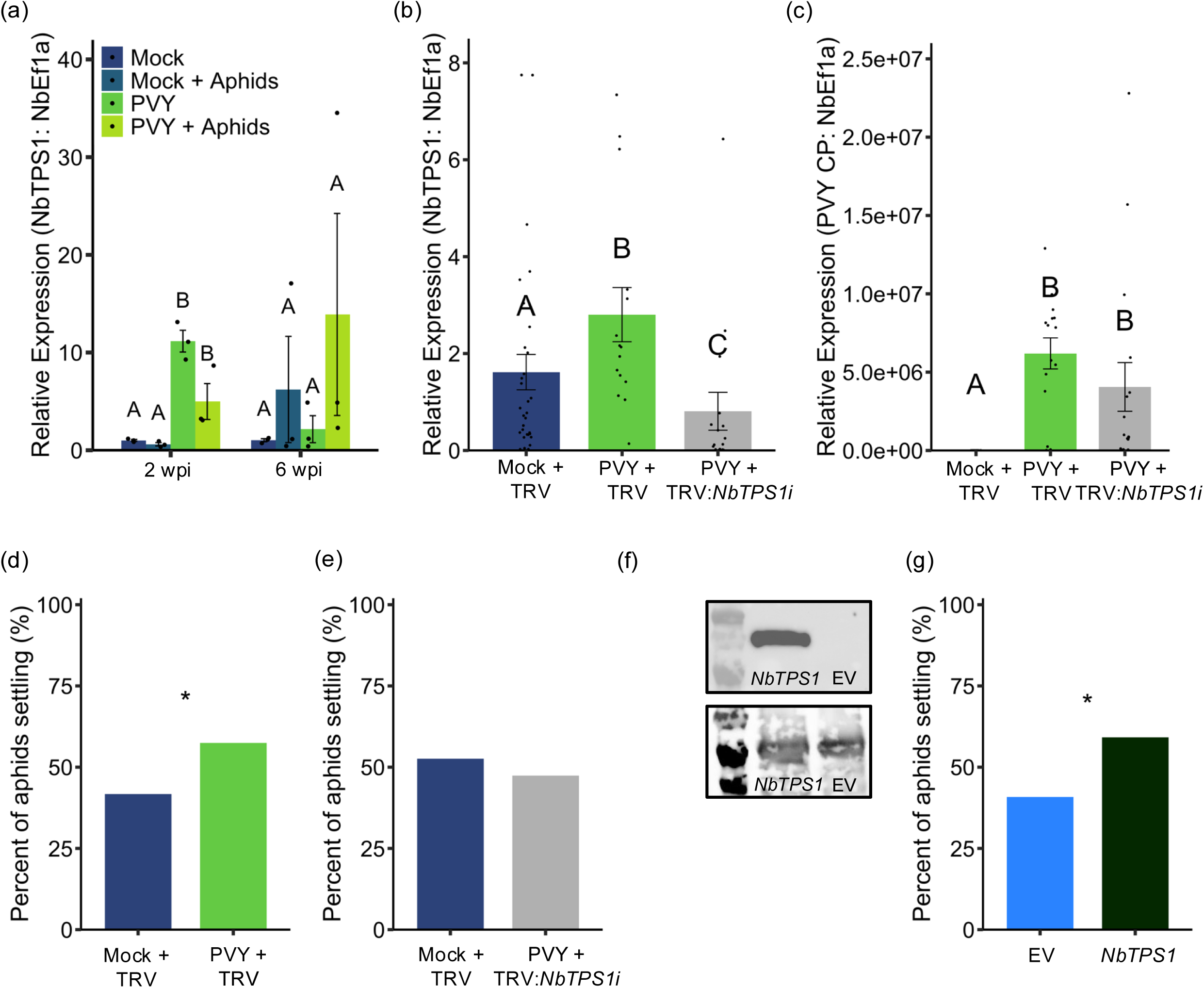
PVY induction of terpene synthase 1 (*NbTPS1*) early in infection is required for increased aphid settlement. (a) Relative expression of terpene synthase 1 (*NbTPS1*) in response to PVY infection at 2- and 6-weeks post-inoculation (wpi) with 48 hours of *Myzus persicae* aphid herbivory. (b) Relative expression of *NbTPS1* in mock-inoculated plants infected with tobacco rattle virus (TRV), PVY-infected plants co-infected with TRV, and PVY-infected plants co-infected with TRV silencing construct (TRV:*NbTPS1*i). (c) Relative expression of PVY coat protein (CP) in mock-inoculated plants infected with TRV, PVY-infected plants co-infected with TRV, and PVY-infected plants co-infected with TRV:*NbTPS1i.* Adult aphids were given a choice between (d) mock-inoculated and PVY-infected leaves co-infected with TRV or (e) between mock-inoculated leaves co-infected with TRV or PVY-infected leaves co-infected with TRV:*NbTPS1i.* (f) Western blot showing *NbTPS1* protein accumulation in leaves expressing pMDC32 *NbTPS1* but not for the empty expression vector (pMDC32 EV). The lower panel is the ponceau stain showing equal protein loading. (g) Adult aphids were given a choice between leaves transiently overexpressing pMDC32 EV or pMDC32:*NbTPS1.* Bars indicate standard errors of the mean. Letters indicate significant differences of *P* < 0.05 between treatments as identified by TukeyHSD for (a) (N = 3) and to Kruskal-Wallis for (b) and (c) (N = 15-31). Asterisks indicate significant differences of *P* < 0.05 according to a Log-likelihood test of independence for (d), (e), and (f) (N = 13-15). For (a), (b) and (c) transcripts are expressed relative to *NbEf1ɑ*).

To determine if *NbTPS1* expression mediates the attraction of aphids to PVY^-^infected plants at 2 wpi (Fig. 1b), we used a TRV-VIGS system to silence *NbTPS1* expression in PVY-infected plants (Fig. 4b). *NbTPS1* silencing had no significant impact on PVY titer in infected plants (Fig. 4c). When given a choice between mock-inoculated and PVY-infected *N. benthamiana* plants co-infected with a control TRV construct, aphids still preferred to settle on the PVY and TRV co-infected plants compared (Fig. 4d; G = 6.376, *P* = 0.012). However when *NbTPS1* was silenced with TRV in PVY-infected plants (TRV:*NbTPS1i*) aphids no longer preferred to settle on the co-infected plants (Fig. 4e; G = 0.621, *P* = 0.431). To confirm that TRV was not influencing aphid choice, we performed a control choice test between mock-inoculated plants with and without TRV infection and found that there was no difference in aphid settlement (Fig. S2; G = 0.148 *P* = 0.700).

To further investigate the role of *NbTPS1* in aphid preference, we transiently overexpressed *NbTPS1* in leaves of healthy *N. benthamiana* and performed choice tests. Overexpression was confirmed by Western blot (Fig. 4f). Aphid settlement was significantly greater on leaves overexpressing *NbTPS1* compared to leaves expressing an empty vector plasmid (Fig. 4g; G = 9.575, *P* = 0.002). Taken together, these results indicate that PVY induction of *NbTPS1* at 2 wpi mediates aphid attraction to PVY-infected plants at early stages of infection.

## Discussion

In this study, we demonstrate that *M. persicae* aphid vectors have increased settlement and fecundity on PVY-infected *N. benthamiana* plants 2 weeks post inoculation, an effect absent later in infection (Fig. 1a,b). Using RNA-seq and qRT-PCR we show the expression of one sesquiterpene synthase gene, *NbTPS1*, was significantly increased by PVY early in infection, however there was no significant induction by PVY at later time points (Fig. 3b; Fig. 4a). While previous studies in potato and tobacco have shown that PVY upregulates terpene synthase genes (Chen *et al*., 2017; Osmani *et al*., 2019; Ross *et al*., 2022), we go further and demonstrate that PVY temporally modulates the expression of *NbTPS1* to mediate aphid vector attraction using VIG silencing and overexpression of *NbTRS1* (Fig. 4). Modulating *NbTPS1* expression to increase vector attraction to infected plants early in infection but not at later time points may benefit PVY spread by discouraging revisits to already infected plants. Similarly, this strategy would increase the likelihood that viruses are acquired by suitable vector for transmission.

Terpenoids are a large and diverse group of VOCs that mediate plant interactions with insects and other organisms (Paré & Tumlinson, 1999; Tholl, 2006; Cheng *et al*., 2007). *NbTPS1* encodes a variety of terpenoid products including α-bergamotene, α-cedrene, β-cedrene, α-himachalene, α-longipinene, epi-aristolochene, and cedrol (Li *et al*., 2014, 2015). Previous studies have demonstrated a role of *NbTPS1* in mediating plant-virus-vector interactions in begomovirus and whitefly systems (Li *et al*., 2014; Wang *et al*., 2023). Tomato yellow leaf curl China virus (TYLCCNV), for example, attracts whitefly (*Bemisia tabaci*) vectors to *N. benthamiana* by using a betasatellite-encoded βC1 protein. βC1 competes with the MYC2 binding bHLH domain, interfering with MYC dimerization and suppressing two MYC2-regulated TPS genes, including *NbTPS1*. This reduces α-bergamotene emissions and increases whitefly (*Bemisia tabaci*) vector attraction to infected plants (Li *et al*., 2014; Wang *et al*., 2023). In another study it was shown that TYLCCNV infection suppresses the expression of 5-epi-aristolochene synthase in tobacco, reducing α-cedrene and β-cedrene emissions, and increasing whitefly vector population growth on infected plants compared to controls (Luan *et al*., 2013). In contrast to these studies, we found that the potyvirus PVY induces *NbTPS1* expression to increase aphid vector attraction to infected plants (Fig. 4b,d). Previous studies have shown that PVY can both induce and suppress emission of various volatile terpenes in potato (Eigenbrode *et al*., 2002; Petek *et al*., 2014), and it is possible the PVY also alters the expression of *TPS* genes. It is not known how PVY increases *NbTPS1* transcripts. Future experiments on the impacts of PVY on plant-insect interactions should take this into account.

Aphids use visual and chemical cues to locate suitable host plants (de Vos & Jander, 2010; Schröder *et al*., 2017). In this study, we performed all aphid choice tests bioassays in a growth chamber overnight. Therefore, there were periods of time during the assay where the growth chamber lights were on, and we cannot rule out that aphid choice was not influenced by visual symptomatic differences in PVY-infected plants compared to mock-inoculated controls. However, our VIGS studies, where both sets of plants were infected and displayed similar symptoms, suggest that the observed aphid preferences are likely influenced by chemical cues rather than visual cues (Fig. 4d,e). Additionally, overexpressing *NbTPS1* in healthy *N. benthamiana* leaves did not induce visual changes, further supporting the role of chemical cues mediating aphid behavior (Fig. 4g). Future experiments focused on measuring specific changes in *N. benthamiana* volatiles with the various genetic tools we employed could be used to further dissect this.

We found that *Myzus persicae* fecundity was enhanced on PVY-infected plants early in infection, but not later (Fig. 1a) despite higher viral titer at later stage infection (Fig. 1c). This could be due to increased upregulation of defense-related transcriptional pathways later in PVY infected plants later in infection (Fig. 2e,f; C17 and C19). Indeed, viruses have been shown to manipulate plant defenses responses in ways that attract vectors to infected plants and subsequently encourage insect dispersal to enhance virus transmission (Carr *et al*., 2018; Wu & Ye, 2020). Cucumber mosaic virus (CMV) increases *M. persicae* preference to infected *Cucurbita pepo* plants through changes in plant VOC profile but reduces aphid performance on infected plants through reduced host quality compared to control plants (Mauck *et al*., 2010). Similarly, viruses can manipulate jasmonate signaling pathways via disruption of MYC interactions with repressor proteins, which attenuates JA-mediated defenses and enhances vector performance (Wu & Ye, 2020). For example, the non-structural protein (NSs) from tobacco spotted wilt virus (TSWV targets and suppresses MYC family transcription factors increasing the attraction and population growth of Western flower thrip (*Frankiniella occidentalis*) vectors, in both pepper and Arabidopsis (Wu *et al*., 2019). Alternatively, enhanced aphid fecundity on PVY-infected plants early but not later in infection could be the result of aphid ability to suppress of plant defenses in early-stage PVY-infected plants. It is well known that aphid feeding can subvert plant defenses through secretion of effector proteins that manipulate host cellular processes and interfere with defense signaling pathways (Ray & Casteel, 2022). Aphids may have been better able to secrete effectors in younger PVY-infected plants compared to older PVY-infected plants or there may be synergetic effects between PVY-infection and aphid feeding on plant defenses that enhance aphid performance in younger plants.

While significant progress has been made in understanding how plant viruses influence plant-insect interactions at the transcriptional level (Zanardo *et al*., 2019), our understanding of viral regulation of plant VOC-related gene transcription remains limited. Transcriptomic approaches, such as RNA-seq, offer promising opportunities to dissect the underlying molecular mechanisms mediating plant-virus-vector interactions. Future studies incorporating transcriptomic approaches with chemical ecology will result in a more comprehensive understanding of the intricate molecular mechanisms mediating plant-virus-vector interactions. This integrated approach will not only provide a detailed understanding of plant defense systems but will also shed light on the chemical cues and signaling molecules involved in shaping these ecological interactions. Moreover, identifying transcripts that mediate these interactions will provide targets for future gene editing strategies to attenuate viral spread within plant populations.

## Acknowledgements

The authors thank Maria Persuad-Fernandez for help with plant and aphid colony care and Tyseen Murad for construction of the choice test arenas. We also thank Zoe Economos and Dr. Elias Bloom for valuable feedback on previous versions of the manuscript. This research was supported by a USDA NIFA award to CTN (2020-67034-31787) and a US National Science Foundation award to CLC (1723926).

## Author contributions

CTN and CLC conceived the research project. CTN and MD conducted the experiments. CTN, JS, SGM, PG, SR, and CLC analyzed and visualized the data. CTN and CLC wrote the manuscript with contributions from JS, MD, SGM, PG, and SR.

## SUPPORTING FIGURES

**Fig. S1.**
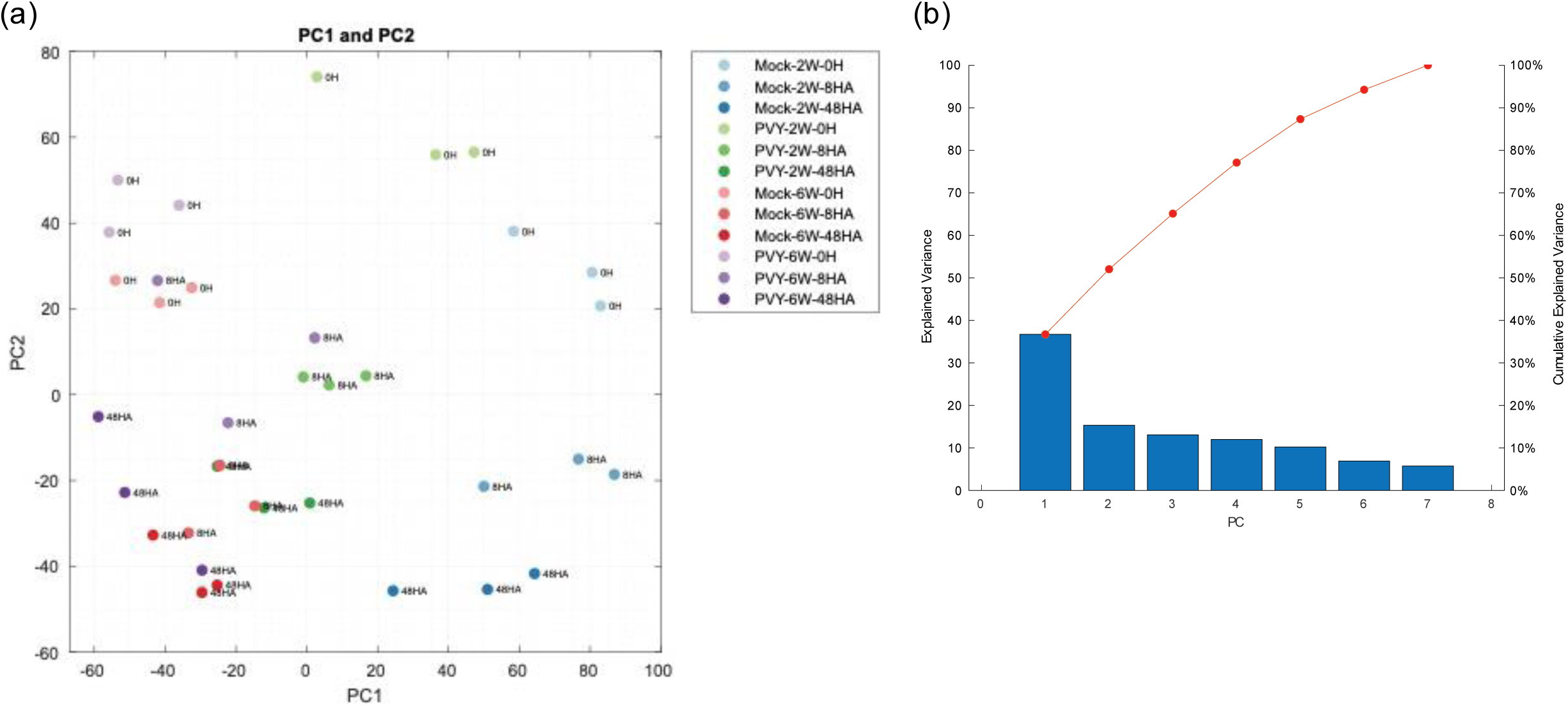
(a) Principal component analysis of triplicate sample from RNA-Seq early- and late-stage (2 wpi, 6 wpi) mock-inoculated and PVY-infected *N. benthamiana* plants without *M. persicae* herbivory (0H) and with 8- and 48 hours of herbivory (8HA, 48HA). (b) The variance explained by each Principal component as derived from Principal component analysis. This shows how much variability in the dataset is captured by each of the Principal component.

**Fig. S2.**
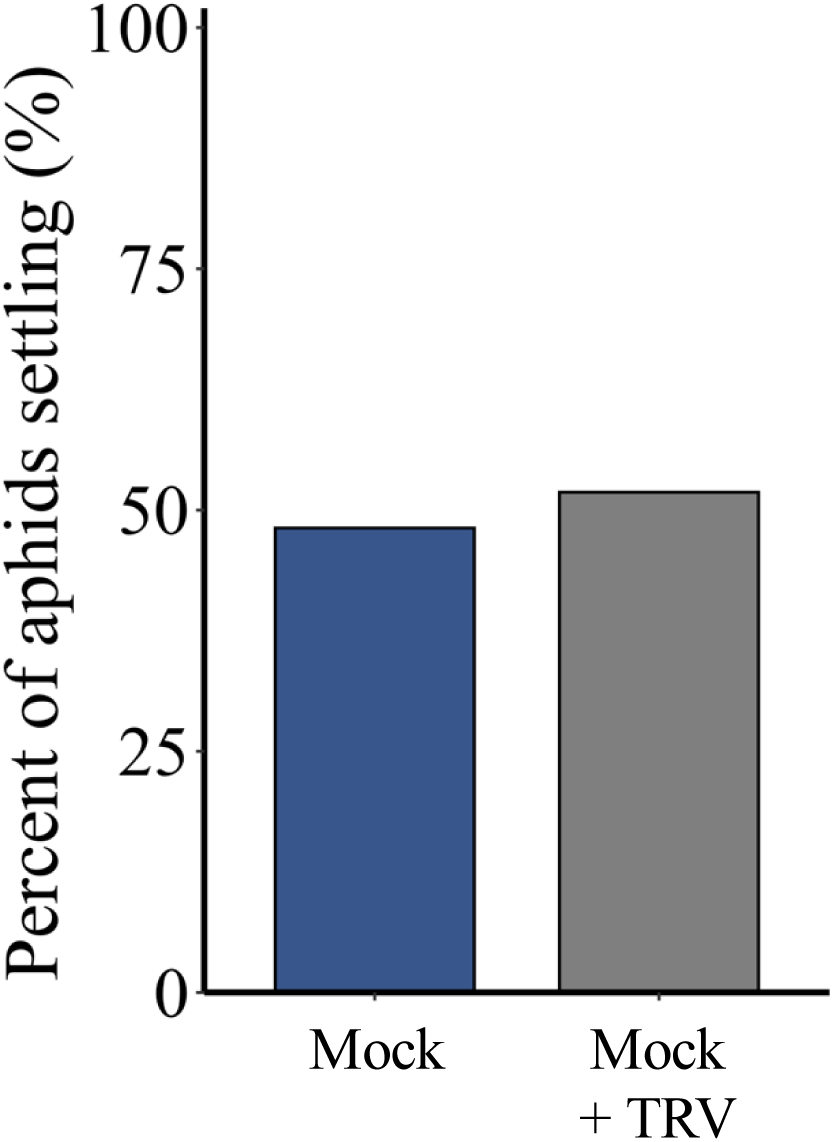
*Myzus persicae* apterous aphids were given a choice between mock-inoculated *N. benthamiana* without (“Mock”) and with co-infection by a tobacco rattle virus empty vector (“Mock + TRV”). There was no difference in aphid attraction to mock-inoculated plants or mock-inoculated plants infected with TRV. Log-likelihood test of independence (N = 6).

